# Altered orbitofrontal sulcogyral patterns in gambling disorder: a multicenter study

**DOI:** 10.1101/439034

**Authors:** Yansong Li, Zixiang Wang, Isabelle Boileau, Jean-Claude Dreher, Sofie Gelskov, Alexander Genauck, Juho Joutsa, Valtteri Kaasinen, José Perales, Nina Romanczuk-Seiferth, Cristian M Ruiz de Lara, Hartwig R Siebner, Ruth J van Holst, Tim van Timmeren, Guillaume Sescousse

## Abstract

Gambling disorder is a serious psychiatric condition characterized by decision-making and reward processing impairments that are associated with dysfunctional brain activity in the orbitofrontal cortex (OFC). However, it remains unclear whether OFC functional abnormalities in gambling disorder are accompanied by structural abnormalities. We addressed this question by examining the organization of sulci and gyri in the OFC. This organization is in place very early and stable across life, such that OFC sulcogyral patterns (classified into Type I, II and III) can be regarded as potential pre-morbid markers of pathological conditions. We gathered structural brain data from nine existing studies, reaching a total of 165 individuals with gambling disorder and 159 healthy controls. Our results, supported by both frequentist and Bayesian statistics, show that the distribution of OFC sulcogyral patterns is skewed in individuals with gambling disorder, with an increased prevalence of Type II pattern compared with healthy controls. Examination of gambling severity did not reveal any significant relationship between OFC sulcogyral patterns and disease severity. Altogether, our results provide evidence for a skewed distribution of OFC sulcogyral patterns in gambling disorder, and suggest that pattern Type II might represent a pre-morbid structural brain marker of the disease. It will be important to investigate more closely the functional implications of these structural abnormalities in future work.

## Introduction

Gambling disorder, from here onwards referred to as pathological gambling^a^, is a behavioral addiction with severe consequences, including bankruptcy, relationship problems and suicide^1^. Consistent with the idea that psychiatric disorders have a biological basis in the brain^2^, functional neuroimaging studies have revealed a core network of dysfunctional brain regions in individuals suffering from pathological gambling^3^. Central to this network is the orbitofrontal cortex (OFC), which displays abnormal activity across a number of cognitive tasks, including expected reward valuation^4, 5^, monetary reward processing^6–8^, risky decision-making^9, 10^ and conflict monitoring^11^. However, it remains unclear whether this alteration in OFC brain function reflects underlying structural abnormalities.

There is a large body of research suggesting that individual variability in brain function is closely related to individual variability in brain structure^12^. Using magnetic resonance imaging, most studies have focused on gray matter volume and cortical thickness as meaningful sources of structural variability. While a few studies have reported decreased OFC gray matter volume^13–15^ and decreased cortical thickness^16^ in pathological gamblers compared with healthy controls, other studies have failed to report significant group differences^17–20^. These inconsistencies might reflect the influence of factors such as age, comorbidities and head motion acting as confounds on structural brain measures^21^, as well as the heterogeneity existing among gamblers, as suggested by a recent study which found decreased OFC gray matter volume specifically in gamblers showing low risk-taking^22^. As a matter of fact, structural abnormalities observed in pathological gamblers are less consistent and of more modest magnitude than those reported in substance addiction^19, 20, 23^. Moreover, it is unclear whether these structural abnormalities represent a pre-morbid marker or are a mere consequence of the disease, as observed in other disorders^24, 25^. One way to address this question is to examine the sulcogyral organization of the OFC.

The organization of sulci and gyri in the brain is governed by the cortical folding which occurs in the perinatal period and leads to sulcogyral patterns that are stable across life^26, 27^. As such, these patterns can be regarded as reliable structural traits that provide an opportunity to investigate possible pre-morbid markers of psychiatric disorders, independently of confounding factors such as illness duration or medication use^28^. In the OFC, sulcogyral patterns have been classified into three different types (Types I, II and III) based on the continuity/discontinuity of the medial and lateral orbitofrontal sulci^29^ (see Fig 1 and Methods for details). In previous work, we have observed that sulcogyral pattern types constrain the location of reward-related value signals in the OFC^30^. In the field of psychiatry, the impact of OFC sulcogyral patterns has been studied extensively in the context of schizophrenia. Nakamura et al.^31^ initially reported a decreased proportion of Type I pattern and an increased proportion of Type II and Type III patterns in individuals suffering from schizophrenia. These findings have been replicated in further studies^32–37^, and extended to individuals at high risk of developing schizophrenia^38, 39^ (but see^40^), suggesting that Types II and III might represent pre-morbid markers of schizophrenia. Among patients with schizophrenia, Type III in particular has been associated with poorer cognitive functioning and IQ, as well as more severe symptoms and impulsivity^31, 37, 40^. However, very little work has been done outside of schizophrenia. While one study reported an increased prevalence of Type III in autism spectrum disorders^41^, another study showed that Type III was associated with greater lifetime cannabis consumption in cannabis users^42^.

**Figure 1.**
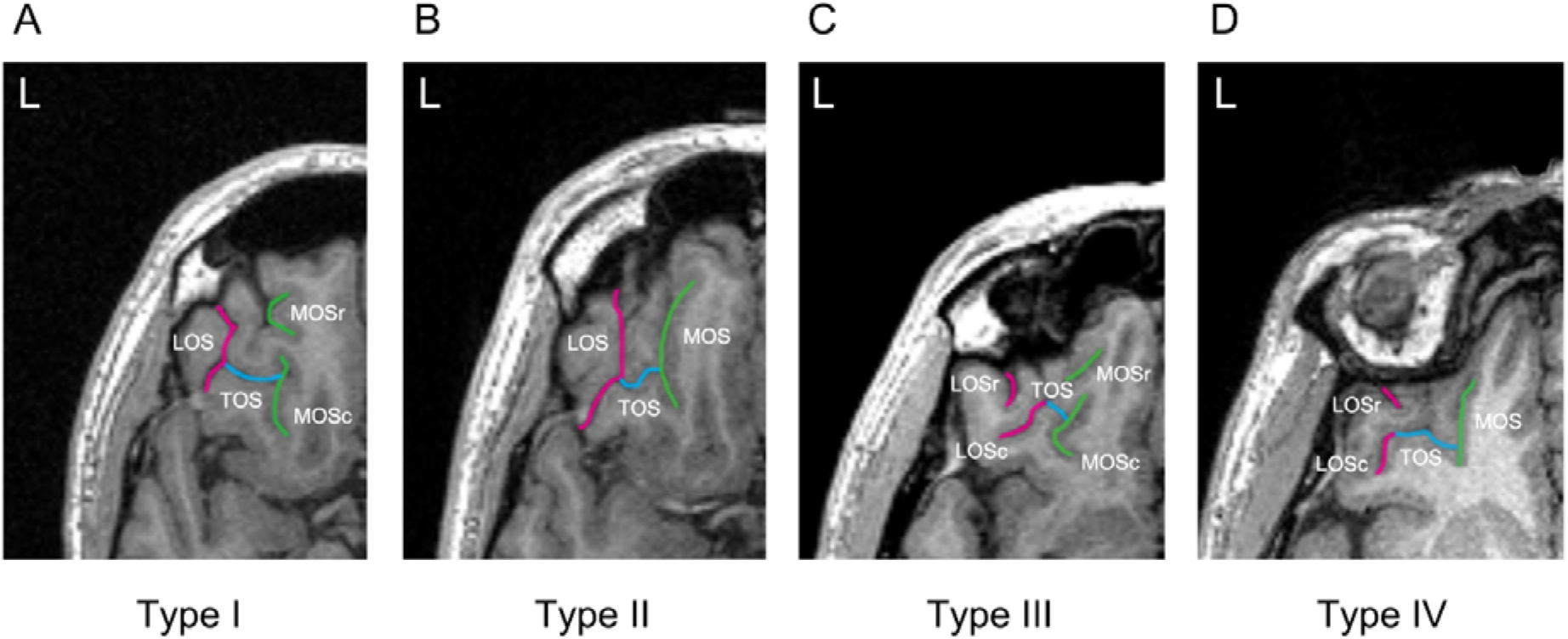
Classification of the orbitofrontal cortex sulcogyral patterns with MRI. Examples of the four major sulcogyral patterns from four different participants. Patterns were classified into four subtypes (Type I – IV) according to the continuity of the LOS and MOS in the rostrocaudal direction (r: rostral, c: caudal). Type I refers to continuous LOS and discontinuous MOS (**A**), Type II refers to continuous LOS and MOS (**B**), Type III refers to discontinuous LOS and MOS (**C**), and Type IV refers to continuous MOS and discontinuous LOS (**D**). Sulcal continuities of the medial and lateral orbital sulci were determined by evaluating several consecutive axial slices rather than just a single slice.

In this study, we aimed to examine the distribution of OFC sulcogyral patterns among pathological gamblers, as well as their relationship with gambling severity, under the premise that the well-described functional impairments reported in the OFC might reflect pre-morbid structural markers. In order to maximize statistical power, we pooled together nine existing structural MRI datasets comprising of 324 individuals. Because we found it hazardous to make predictions based on the existing literature, we refrained from making specific hypotheses and considered this study to be exploratory.

## Materials and Methods

### Participants

A total of 177 pathological gamblers (PGs) and 169 healthy controls (HCs) were included in the present study. These data were pooled from nine separate previous neuroimaging studies^8, 15, 18, 43–48^ (**Supplementary Table 1**). From this aggregated sample, 18 participants (10 PGs and 8 HCs) were excluded because of head movement artefacts on anatomical T1 scans preventing us from reliably identifying OFC sulcogyral patterns, and 4 participants (2 PGs and 2 HCs) were excluded because either demographic or diagnostic information was missing. As a result, 165 PGs (164 men / 1 woman, age = 34.25 ± 10.00 years) and 159 HCs (154 men / 5 women, age = 33.00 ± 9.76 years) were included in the final analysis. The two groups were matched on age, gender and handedness within individual studies, which was verified at the whole-population level (**Table 1**). In addition, all individual studies matched the groups on IQ and/or education level, while 7 out of 9 studies matched the groups on the number of smokers. All participants gave written informed consent to be part of the original studies, which were approved by the local ethics committees.

**Table 1.** Demographic and clinical characteristics of the sample (pooled across 9 studies)

|  | **PGs**<br>**(N=165)** | **HCS**<br>**(N=159)** | **Group comparison** |
| --- | --- | --- | --- |
| Age | 34.25 ± 10.00 | 33.00 ± 9.76 | $t(322)=1.14, p=0.257$ |
| Gender (M/F) | 164/1 | 154/5 | $\chi^2=1.64, p=0.200$ |
| Handedness(R/L/mixed) | 152/11/2 | 149/4/6 | $p=0.080$ (FET) |
| SOGS | 10.30 ± 4.14 (N=148) | 0.41 ± 0.88 (N=144) | $t(160.64)=28.42, p<0.001$ |
| PG-YBOCS | 24.35 ± 7.06 (N=17) | 10.87 ± 1.69 (N=15) | $t(18.05)=7.63, p<0.001$ |

FET = Fisher's exact test; SOGS = South Oaks Gambling Screen; PG-YBOCS = Yale Brown Obsessive Compulsive Scale adapted for Pathological Gambling; PGs = pathological gamblers; HCs = healthy controls. For age, SOGS and PG-YBOCS, numbers represent mean ± standard deviation.

The PGs of all nine studies were diagnosed using psychiatric interviews or questionnaires based on DSM-IV criteria. Furthermore, eight of these studies used the South Oaks Gambling Screen questionnaire (SOGS)^49^ to assess the severity of gambling symptoms, while the last one used the Yale Brown Obsessive Compulsive Scale adapted for Pathological Gambling (PG-YBOCS)^50^ (**Table 1**). None of the HCs had a known history of neurological disorder or current psychiatric Axis I disorder, except for one of them meeting past criteria for alcohol abuse. In the gambling group, given the high co-morbidity between pathological gambling and other psychiatric disorders^51^, gamblers with the following co-morbidities were included: current cannabis dependence (N = 1); past cannabis dependence (N = 1); past cannabis abuse (N = 1); past alcohol dependence (N = 2); past alcohol abuse (N = 1); lifetime history of dysthymia (N = 1); remitted post-traumatic stress disorder (N = 2). In addition, three gamblers used cannabis weekly in the past 6 months before scanning, but did not meet the DSM-IV criteria for abuse/dependence.

### MRI acquisition

T1-weighted structural MR images were independently acquired at each imaging site. Data acquisition details are summarized in **Supplementary Table 1** based on the descriptions from the original studies.

### OFC sulcogyral pattern classification

The procedures for classifying the OFC sulcogyral patterns in this study were based on one of our previous studies^30^. Specifically, the OFC sulcogyral patterns were identified separately in each hemisphere using the medical image analysis software MRIcro (www.mccauslandcenter.sc.edu/mricro/) and classified according to the criteria described by Chiavaras and Petrides^29^. With regard to the orbitofrontal sulci in the human brain, four main sulci have been identified, namely, the olfactory, medial, lateral, and transverse orbital sulci. On the basis of the continuity of the medial and lateral orbital sulci (MOS and LOS, respectively), the original work by Chiavaras and Petrides^29^ classified the morphology of the human orbitofrontal sulci into three main types in each hemisphere (Type I, II, and III), while a fourth type (Type IV) was later identified in a number of studies^36, 38, 40, 41^ (**Fig 1**). In Type I, rostral and caudal portions of the LOS (LOSr and LOSc) are connected to one another, whereas the rostral and caudal portions of the MOS (MOSr and MOSc) are clearly separate (**Fig 1A**). Compared with the Type I pattern, the distinctive feature of the Type II is that rostral portions of both LOS and MOS are connected to their caudal portions, forming the continuous MOS and LOS, and both sulci are jointed by the horizontally oriented transverse orbital sulcus (**Fig 1B**). In Type III, the critical distinctive characteristic is that the rostral and caudal parts of both MOS and LOS are clearly disconnected (**Fig 1C**). In Type IV, the rostral and caudal portions of the LOS were interrupted in the presence of a continuous MOS, thus representing the opposite of Type I (**Fig 1D**). The sulcus continuity was determined by evaluating several adjacent axial slices rather than focusing on one single slice.

Two raters (Y.L. and Z.W.), who were blind to the participants’ identity, independently performed the OFC sulcogyral pattern classification for all participants. Inter-rater reliability (Cohen’s kappa) was 0.84 for the left hemisphere and 0.83 for the right hemisphere. All ambiguous classifications identified in the current sample (i.e. 9% of the total sample) were reviewed by Y.L. and consensus was reached.

### Statistical analyses

All analyses were performed using SPSS for frequentist statistics (http://www.spss.com/software) and JASP for Bayesian statistics (https://jasp-stats.org). Independent-samples two-sided t-tests (with correction for inhomogeneity of variance where appropriate) were performed to assess group differences in age, SOGS and PG-YBOCS. Pearson’s χ^2^ test, or Fisher’ s exact test (when more than 20% of cells had expected counts less than 5) were employed to evaluate group differences in gender and handedness.

Group differences in OFC sulcogyral patterns were evaluated using frequentist χ^2^ tests (producing p-values) and Bayesian contingency table analyses (producing Bayes Factors, BF) on the distribution of Type I, Type II and Type III patterns. Because hemispheres with a Type IV pattern were rare (4% of all hemispheres), they were excluded from the statistical analyses, in line with the procedure used in a number of previous studies^36, 38, 40, 41^. These analyses were performed across the left and right hemispheres, as well as separately in each hemisphere. To further identify which OFC sulcogyral types have a skewed frequency in PGs compared with HCs, post-hoc χ^2^ tests were performed for each sulcogyral pattern separately. Since the latter pairwise analysis involves three comparisons (Type I, II, and III), so we used a Bonferroni-corrected threshold of 0.05/3 = 0.017. In order to verify that the OFC sulcogyral pattern distribution in HCs was comparable to that reported in the reference study of Chiavaras and Petrides^29^, we also compared these distributions statistically using the same tests as above. Finally, we used a categorical regression analysis to examine whether sulcogyral pattern types were associated with clinical symptoms (SOGS scores) in PGs. This latter analysis was performed on N=148 PGs (instead of the full sample N=165 PGs) since one of the studies^45^ did not report SOGS scores. Following the approach used in previous studies^31, 41, 52^, we defined three categorical predictors for the three main sulcogyral patterns (Type I, II and III). For each predictor and each participant, we assigned a value of 2 if the sulcogyral pattern under consideration was present in either hemisphere (i.e. in the left, right, or both hemispheres), and a value of 1 if the sulcogyral pattern was absent in either hemisphere. We report the overall fit as well as the standardized coefficients for each predictor.

## Results

### Group differences in orbitofrontal sulcogyral patterns

**Table 2** and **Figure 2** illustrate the OFC sulcogyral pattern distribution observed in each group (detailed distributions for each study are provided in **Supplementary Tables 2 and 3**). Importantly, the sulcogyral pattern distribution observed in HCs was very close to that reported in the original study of Chiavaras and Petrides^29^ (Left hemisphere: *χ*^*2*^ = 0.02, *p* = 0.991, *BF*_*10*_ = 0.139; Right hemisphere: *χ*^*2*^ = 2.25, *p* = 0.324, *BF*_*10*_ = 0.347; All hemispheres: *χ*^*2*^ = 0.62, *p* = 0.734, *BF*_*10*_ = 0.173). This indicates that any group difference in the present study is more likely to result from an unusual distribution in the gambling group rather than in the healthy group. In addition, and in line with the results from Nakamura et al.^31^, we did not find evidence supporting a relationship between the left and the right hemispheres in terms of type of OFC sulcogyral pattern, as revealed by likelihood ratio (LR) tests in multinomial logistic regressions (HCs: *LRχ*^*2*^ = 3.32, *p* = 0.51; PGs: *LRχ*^*2*^ = 8.81, *p* = 0.08; detailed distributions of OFC sulcogyral patterns as a function of left-right combination are reported in **Supplementary Table 4**). We thus treated hemispheres as independent and pooled them together in some of the subsequent analyses.

**Table 2.**
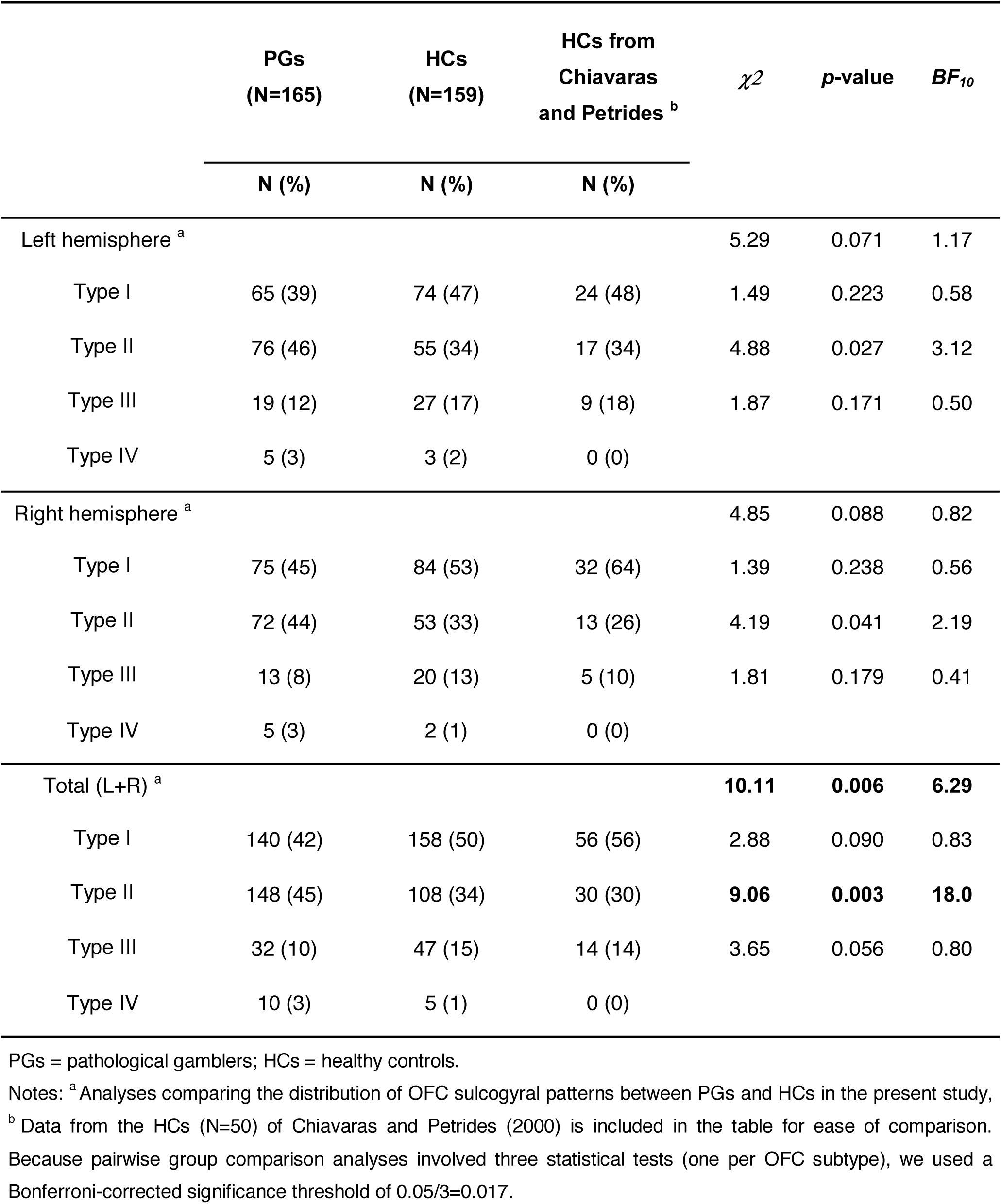
Distribution of OFC sulcogyral patterns.

| | PGs<br>(N=165) | HCs<br>(N=159) | HCs from<br>Chiavaras<br>and Petrides <sup>b</sup> | $\chi^2$ | *p*-value | *BF*<sub>10</sub> |
| --- | --- | --- | --- | --- | --- | --- |
|  | N (%) | N (%) | N (%) |  |  |  |
| Left hemisphere <sup>a</sup> |  |  |  | 5.29 | 0.071 | 1.17 |
| Type I | 65 (39) | 74 (47) | 24 (48) | 1.49 | 0.223 | 0.58 |
| Type II | 76 (46) | 55 (34) | 17 (34) | 4.88 | 0.027 | 3.12 |
| Type III | 19 (12) | 27 (17) | 9 (18) | 1.87 | 0.171 | 0.50 |
| Type IV | 5 (3) | 3 (2) | 0 (0) |  |  |  |
| Right hemisphere <sup>a</sup> |  |  |  | 4.85 | 0.088 | 0.82 |
| Type I | 75 (45) | 84 (53) | 32 (64) | 1.39 | 0.238 | 0.56 |
| Type II | 72 (44) | 53 (33) | 13 (26) | 4.19 | 0.041 | 2.19 |
| Type III | 13 (8) | 20 (13) | 5 (10) | 1.81 | 0.179 | 0.41 |
| Type IV | 5 (3) | 2 (1) | 0 (0) |  |  |  |
| Total (L+R) <sup>a</sup> |  |  |  | **10.11** | **0.006** | **6.29** |
| Type I | 140 (42) | 158 (50) | 56 (56) | 2.88 | 0.090 | 0.83 |
| Type II | 148 (45) | 108 (34) | 30 (30) | **9.06** | **0.003** | **18.0** |
| Type III | 32 (10) | 47 (15) | 14 (14) | 3.65 | 0.056 | 0.80 |
| Type IV | 10 (3) | 5 (1) | 0 (0) |  |  |  |

PGs = pathological gamblers; HCs = healthy controls.
Notes: <sup>a</sup> Analyses comparing the distribution of OFC sulcogyral patterns between PGs and HCs in the present study,
<sup>b</sup> Data from the HCs (N=50) of Chiavaras and Petrides (2000) is included in the table for ease of comparison. Because pairwise group comparison analyses involved three statistical tests (one per OFC subtype), we used a Bonferroni-corrected significance threshold of 0.05/3=0.017.

**Figure 2.**
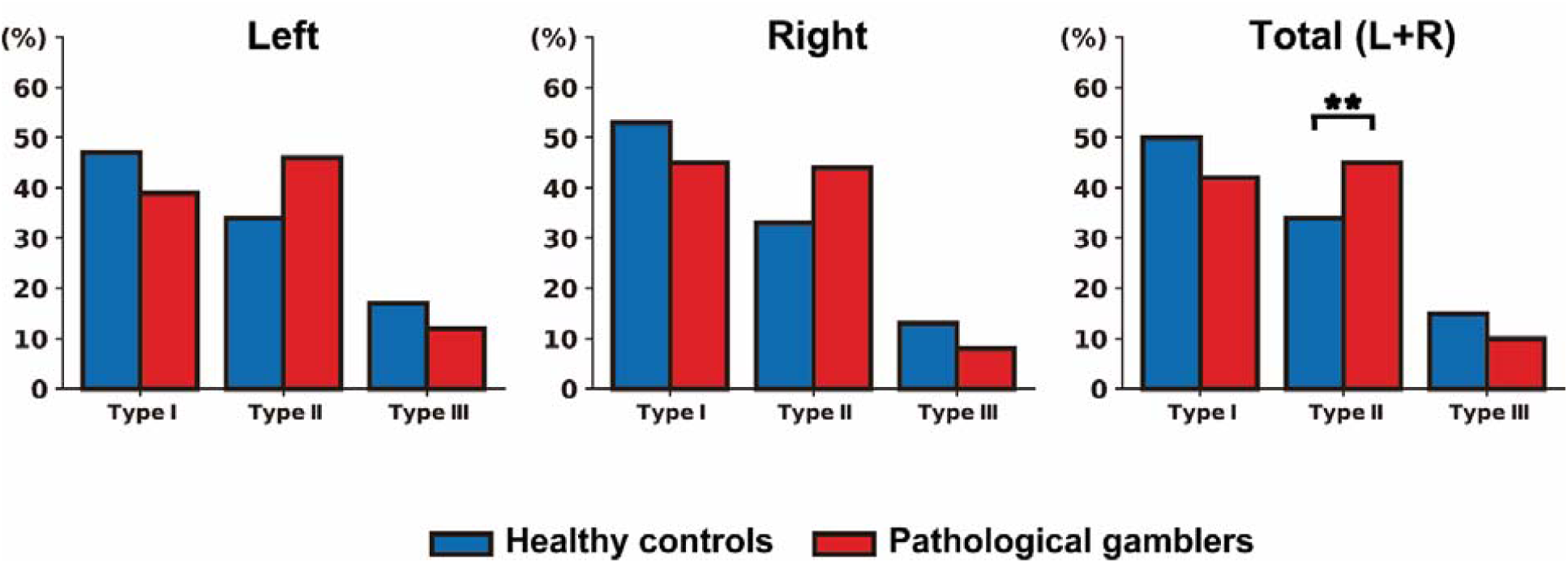
Distribution of the orbitofrontal cortex sulcogyral patterns in the left hemisphere, right hemisphere and across both hemispheres, in both pathological gamblers (N = 165) and healthy controls (N = 159). **denotes p < 0.005.

Across the left and right hemispheres, a χ^2^ analysis revealed that the OFC sulcogyral pattern distribution in PGs was significantly different from that observed in HCs (*χ*^*2*^ = 10.11, *Cramer’s V* = 0.126, *p* = 0.006). In order to quantify the evidence in favor of a group difference, we performed a Bayesian contingency table analysis based on a Poisson sampling model^53^, using the default uniform priors implemented in JASP. The resulting Bayes Factor (*BF*_*10*_ = 6.29) indicated that our data were about 6 times more likely under the hypothesis of a group difference than under the null hypothesis of no group difference. Such a Bayes Factor is traditionally interpreted as moderate evidence for the alternative hypothesis that there is indeed an association between the group and OFC sulcogyral pattern distribution. Post-hoc tests further showed that the group difference was primarily driven by a significantly enhanced prevalence of Type II pattern in PGs compared with HCs (45% vs 34%, *χ*^*2*^ = 9.06, *Cramer’s V* = 0.120, *p* = 0.003). The associated Bayes Factor (*BF*_*10*_ = 18.0) confirmed this result, indicating strong evidence for a different proportion of Type II pattern between PGs and HCs. In contrast, we found no significant difference in the distribution of Type I (*χ*^*2*^ = 2.88, *p* = 0.090, *BF*_*10*_ = 0.83) and Type III patterns (*χ*^*2*^ = 3.65, *p* = 0.056, *BF*_*10*_ = 0.80). Also, when examining the left and right hemispheres separately, the group difference in OFC sulcogyral pattern distribution did not reach significance (left: *χ*^*2*^ = 5.29, *p* = 0.071, *BF*_*10*_ = 1.17: right: *χ*^*2*^ = 4.85, *p* = 0.088, *BF*_*10*_ = 0.82). Note though that the distribution of OFC sulcogyral patterns was very similar between the left and right hemispheres in either HCs (*χ*^*2*^ = 1.71, *p* = 0.425, *BF*_*10*_ = 0.20) or PGs (*χ*^*2*^ = 1.95, *p* = 0.378, *BF*_*10*_ = 0.19).

With regards to left/right hemisphere combinations, 22.4% of PGs (compared with 12.6% of HCs) had a Type II pattern in both hemispheres (*χ*^*2*^ = 5.41, *Cramer’s V* = 0.129, *p* = 0.020; *BF*_*10*_ = 3.12). Furthermore, 67.3% of PGs (compared with 55.3% of HCs) had at least one hemisphere with a Type II pattern (*χ*^*2*^ = 4.86, *Cramer’s V* = 0.122, *p* = 0.027; *BF*_*10*_ = 3.01). In terms of relative risk, participants with a Type II pattern in both hemispheres had a 1.35-fold increased risk of being in the gambling group, compared with participants who had only one or zero Type II pattern in either hemisphere. Similarly, participants with at least one hemisphere with a Type II pattern had a 1.29-fold increased risk of being in the gambling group, compared with participants without any Type II pattern.

### OFC sulcogyral patterns and clinical measures

The categorical regression analysis revealed no significant relationship between OFC sulcogyral pattern types and total SOGS scores in PGs (overall fit: *F*(3,144) = 0.10, *p* = 0.963 - standardized coefficients: *β*_Type I_ = -0.03, *F* = 0.08, *p* = 0.779; *β*_Type II_ = −0.05, *F* = 0.27, *p* = 0.600; *β*_Type III_ = −0.02, *F* = 0.06, *p* = 0.810).

## Discussion

This study is the first to investigate the distribution of OFC sulcogyral patterns in pathological gambling, and more broadly, in addiction. Our results, supported by both frequentist and Bayesian statistics, show that the distribution of OFC sulcogyral patterns is skewed in pathological gamblers, with the Type II pattern showing a moderate increase in prevalence compared with healthy controls. The presence of a Type II pattern was not associated with higher gambling severity. Altogether, these results suggest that Type II pattern might represent a pre-morbid structural brain marker of pathological gambling, but without predictive value for symptom severity.

The finding of a skewed distribution of OFC sulcogyral patterns in pathological gambling is of potential clinical relevance, especially given the lack of reliable neuroanatomical markers for this pathology. It also strengthens the existing evidence supporting a central role for the OFC in pathological gambling^3^. Given that sulcogyral patterns are in place very early during brain development and are stable across the lifespan^26, 27^, our results also support the idea that pathological gambling has a partly neurodevelopmental origin. This idea resonates with the results of previous studies on substance abuse, showing that pre-existing structural traits in the OFC have predictive value for later substance use in adolescents^54, 55^. In particular, Kühn et al.^56^ have shown that lower gyrification in the OFC, presumably related to pre-natal alcohol exposure, was associated with increased alcohol-related problems in the next two years. Similarly, Chye et al.^42^ have found that cannabis users with OFC sulcogyral pattern Type III consumed more cannabis over their lifetime. Together, these results support the hypothesis that neurodevelopmental structural traits in the OFC might predispose to addictive behaviors.

From a mechanistic perspective, the key question is how individual differences in OFC sulcogyral patterns would influence the vulnerability to develop (gambling) addictive behaviors. One idea is that this may be mediated through individual differences in OFC connectivity. Indeed, cortical folding patterns are constrained by white matter tracts, such as the uncinate fasciculus which connects the OFC to the anterior temporal lobe within the limbic system^57^, and are thus thought to reflect structural^58, 59^ as well as functional connectivity^60^. Thus, the skewed distribution of OFC sulcogyral patterns in pathological gamblers may reflect altered connectivity patterns, that may in turn affect important cognitive function of the OFC such as decision-making or behavioral inhibition. Altered OFC sulcogyral patterns and associated changes in OFC connectivity might also be associated with certain personality traits known to predispose to addictive behaviors. In particular, Type II pattern has been associated with positive emotionality^31^ and increased physical anhedonia^61^ in healthy individuals. These seemingly opposite personality traits are key defining features of two sub-types of gamblers known as “antisocial-impulsive” and “emotionally vulnerable”, respectively^62^. Thus, it could be that the increased prevalence of OFC sulcogyral Type II pattern among pathological gamblers in the present study reflects the increased prevalence of positive emotionality and anhedonic traits among these two sub-populations. Interestingly, in healthy individuals, Type III pattern has also been associated with improved regulatory control^31^, a trait known to be protective against addictive behaviors. This would be consistent with a trend towards a lower prevalence of Type III pattern among gamblers compared with healthy controls in the present study (although the differences did not reach significance). Future studies should explicitly test these hypotheses.

This study has various strengths. The sample size was larger than in any previous structural imaging study on pathological gambling, and was larger than in most previous studies investigating OFC sulcogyral patterns in psychiatric disorders. The OFC sulcogyral pattern classification was performed independently by two raters for all participants, thus reducing the chances of misclassification. We also used a combination of Frequentist and Bayesian statistics, and applied stringent Bonferroni correction for multiple comparisons where appropriate.

This study also has some limitations. First, the interpretation of our structural brain results in terms of functional consequences is limited by the fact that we did not have functional assessments at hand, such as behavioral scores, questionnaires or brain activity measures. In particular, we were not able to distinguish between different sub-types of gamblers, while these sub-types may be associated with different brain abnormalities as suggested above. This is the main drawback of our strategy to increase statistical power by pooling together datasets from different studies, since the functional assessments did not overlap between these studies. Future work needs to elucidate the link between OFC sulcogyral patterns and their functional implications. Furthermore, the effect sizes we found are relatively small (Cramer’s V between 0.12 and 0.13), suggesting that the observed increased prevalence of Type II pattern is a relatively subtle effect that only affects a minority of gamblers. This might explain why the group difference in OFC sulcogyral pattern type distribution did not reach significance when examined in each hemisphere separately, even though these distributions were qualitatively very similar to the one seen across both hemispheres combined. Finally, we did not find a relationship between OFC sulcogyral patterns and gambling severity. While this might sound surprising, several studies that have reported skewed distribution of OFC sulcogyral patterns in schizophrenia similarly did not observe significant relationships with symptom severity^32, 34^. As suggested by these studies, it could be that OFC sulcogyral patterns do not affect disease progression, or only later in the course of the disease.

In conclusion, our results provide evidence for a skewed distribution of OFC sulcogyral patterns in pathological gambling in a large cohort of individuals studied across different neuroimaging centers. Since this was an exploratory study, it will be very important to replicate this result in an independent sample. Also, based on previous reports of gender differences in the distribution of OFC sulcogyral patterns in other disorders such as schizophrenia^35, 37^, it will be important to test whether our results –observed in an almost entirely male sample– are generalizable to a female gambling population. Finally, future work should examine whether the increased prevalence of Type II pattern observed in pathological gambling extends to substance addiction. Given the similarities between pathological gambling and substance addiction, especially in terms of OFC dysfunction^63^, one could expect a similarly skewed distribution of OFC sulcogyral patterns across the two disorders, in line with the idea of a general vulnerability factor that would predispose to addictive behaviors. However, recent work has also suggested that the neurobiology of pathological gambling and substance addiction might be more different than previously thought, in particular in terms of brain structure alterations^64^, and it could be that the present results are specific to pathological gambling. Ultimately, it will be crucial to design transdiagnostic studies jointly assessing structural and functional alterations in the OFC, in order to better understand how these two levels of alteration relate to each other, and how much they generalize across disorders.

## Acknowledgments

Yansong Li was supported by the National Natural Science Foundation of China (Grant No. 31600929) and the Fundamental Research Funds for the Central Universities (010914380002). Guillaume Sescousse was supported by a Veni grant from the Netherlands Organization for Scientific Research (Grant No. 016.155.218). Juho Joutsa was supported by the Academy of Finland (Grant No. 295580), the Finnish Medical Foundation, and the Finnish Foundation for Alcohol Studies. Valtteri Kaasinen was supported by the Academy of Finland (Grant No. 256836) and the Finnish Foundation for Alcohol Studies. Sofie Gelskov and Hartwig R Siebner were supported by the Danish Council for Independent Research in Social Sciences through a grant to Thomas Ramsøy (“Decision Neuroscience Project”; Grant No. 0601-01361B) and by the Lundbeck Foundation through a Grant of Exellence to Hartwig R Siebner (“ContAct”; Grant No. R59 A5399). Alexander Genauck was supported by Deutsche Forschungsgemeinschaft (DFG) HE2597/15–1, HE2597/15-2, and DFG Graduiertenkolleg 1589/2 “Sensory Computation in Neural Systems”. Nina Romanczuk-Seiferth was supported by a research grant by the Senatsverwaltung für Gesundheit und Soziales, Berlin, Germany (Grant No. 002-2008/ I B 35). Cristian M. Ruiz de Lara and José C. Perales were supported by a grant from the Spanish Government (Ministerio de Economía y Competitividad, Secretaría de Estado de Investigación, Desarrollo e Innovación; Convocatoria 2017 de Proyectos I+D de Excelencia, Spain; co-funded by the Fondo Europeo de Desarrollo Regional, FEDER, European Union; Grant No. PSI2017-85488-P). Jean-Claude Dreher was supported by “LABEX ANR-11-LABEX-0042” of Université de Lyon within the program Investissements d’Avenir (ANR-11-IDEX-007) operated by the French National Research Agency and by a grant from the Fondation pour la Recherche Médicale (Grant No. DPA20140629796).

## Author contributions

Y.L. and G.S. conceived and designed the study; Y.L. and Z.W. analyzed the data; Y.L., Z.W. and G.S. wrote the manuscript; I.B., JCD, SG, AG, JJ, VK, JP, NRS, CRL, HRS, RJvH and TvT edited the manuscript. IB, JCD, SG, AG, JJ, VK, JP, NRS, CRL, HRS, RJvH and TvT are listed alphabetically.

## Footnotes

a Here we prefer to employ the word “pathological gambling”, since all participants were recruited based on the diagnostic criteria of the DSM-IV, i.e. before it was renamed “gambling disorder”.

