## Supplementary Materials for "Altered orbitofrontal sulcogyral patterns in gambling disorder: a multicenter study"

**Supplementary Table 1.** Information on individual studies.

|  | *Boileau et al., 2014* | *Gelskov et al., 2016* | *Genauck et al., 2018* | *Joutsa et al., 2011* | *Ruiz de Lara et al., 2018* | *Romanczuk-Seiferth et al., 2015* | *Sescousse et al., 2013* | *Sescousse et al., 2016* | *van Timmeren et al., 2017* |
| --- | --- | --- | --- | --- | --- | --- | --- | --- | --- |
| Number of pathological gamblers (Number excluded) | 13 (1) | 15 (0) | 22 (4) | 12 (0) | 26 (5) | 19 (2) | 20 (0) | 22 (0) | 28 (0) |
| Number of healthy controls (Number excluded) | 11 (1) | 15 (0) | 23 (1) | 12 (2) | 27 (2) | 17 (2) | 19 (1) | 22 (0) | 23 (1) |
| Manufacturer and Model of scanner | GE Signa | Siemens  Magnetom Tim Trio | Siemens  Magnetom Tim Trio | Philips Gyro-  scan Intera | Siemens Magnetom Tim Trio Syngo MR B17 | Siemens Magnetom Tim Trio | Siemens Sonata | Siemens Magnetom Tim Trio | Philips Intera |
| Field strength (Tesla) | 1.5 | 3 | 3 | 1.5 | 3 | 3 | 1.5 | 3 | 3 |
| Number of channels of head coil | 8 | 8 | 12 | 8 | 32 | 12 | 8 | 32 | 8 |
| Repetition Time (ms) | 8.9–12 | 1540 | 1900 | 25 | 2300 | 1570 | 1970 | 2300 | 1530 |
| Echo Time (ms) | 5.3–15 | 3.9 | 2.52 | 4.6 | 3.1 | 2.7 | 3.93 | 3.03 | 4.2 |
| Sequence | T1-weighted spoiled gradient recalled acquisition | T1-weighted  MP-RAGE | T1-weighted  MP-RAGE | T1-weighted fast field echo | T1-weighted MP-RAGE | T1-weighted MP-RAGE | T1-weighted MP-RAGE | T1-weighted MP-RAGE | T1-weighted  MP-RAGE |

**Supplementary Table 2.** Orbitofrontal Cortex (OFC) sulcogyral pattern distribution in pathological gamblers by studies.

|  | *Boileau et al., 2014* | *Gelskov et al., 2016* | *Genauck et al., 2018* | *Joutsa et al., 2011* | *Ruiz de Lara et al., 2018* | *Romanczuk-Seiferth et al., 2015* | *Sescousse et al., 2013* | *Sescousse et al., 2016* | *van Timmeren et al., 2017* |
| --- | --- | --- | --- | --- | --- | --- | --- | --- | --- |
| Left OFC sulcogyral pattern, n (%) | | | | | | | | | |
| Type I | 5 (42) | 8 (53) | 8 (44) | 4 (33) | 8 (38) | 8 (47) | 5 (25) | 8 (36) | 11 (39) |
| Type II | 5 (42) | 6 (40) | 9 (50) | 6 (50) | 9 (43) | 5 (29) | 11 (55) | 12 (55) | 13 (46) |
| Type III | 1 (8) | 1 (7) | 1 (6) | 2 (17) | 3 (14) | 4 (24) | 4 (20) | 2 (9) | 1 (4) |
| Type IV | 1 (8) | 0 (0) | 0 (0) | 0 (0) | 1 (5) | 0 (0) | 0 (0) | 0 (0) | 3 (11) |
| Right OFC sulcogyral pattern, n (%) | | | | | | | | | |
| Type I | 5 (42) | 7 (47) | 7 (38) | 3 (25) | 11 (53) | 10 (59) | 10 (50) | 8 (36) | 14 (50) |
| Type II | 4 (33) | 7 (47) | 10 (56) | 7 (59) | 7 (33) | 5 (29) | 10 (50) | 9 (41) | 13 (46) |
| Type III | 2 (17) | 0 (0) | 0 (0) | 1 (8) | 3 (14) | 2 (12) | 0 (0) | 4 (19) | 1 (4) |
| Type IV | 1 (8) | 1 (6) | 1 (6) | 1 (8) | 0 (0) | 0 (0) | 0 (0) | 1 (4) | 0 (0) |

**Supplementary Table 3.** Orbitofrontal Cortex (OFC) sulcogyral pattern distribution in healthy controls by studies.

|  | *Boileau et al., 2014* | *Gelskov et al., 2016* | *Genauck et al., 2018* | *Joutsa et al., 2011* | *Ruiz de Lara et al., 2018* | *Romanczuk-Seiferth et al., 2015* | *Sescousse et al., 2013* | *Sescousse et al., 2016* | *van Timmeren et al., 2017* |
| --- | --- | --- | --- | --- | --- | --- | --- | --- | --- |
| Left OFC sulcogyral pattern, n (%) | | | | | | | | | |
| Type I | 4 (40) | 7 (47) | 11 (50) | 5 (50) | 14 (56) | 7 (47) | 7 (39) | 12 (55) | 7 (32) |
| Type II | 4 (40) | 7 (47) | 7 (32) | 3 (30) | 8 (32) | 5 (33) | 7 (39) | 6 (27) | 8 (36) |
| Type III | 2 (20) | 1 (6) | 4 (18) | 2 (20) | 2 (8) | 3 (20) | 4 (22) | 4 (18) | 5 (23) |
| Type IV | 0 (0) | 0 (0) | 0 (0) | 0 (0) | 1 (4) | 0 (0) | 0 (0) | 0 (0) | 2 (9) |
| Right OFC sulcogyral pattern, n (%) | | | | | | | | | |
| Type I | 5 (50) | 7 (47) | 13 (59) | 5 (50) | 11 (44) | 8 (53) | 11 (61) | 11 (50) | 13 (59) |
| Type II | 3 (30) | 8 (53) | 5 (23) | 4 (40) | 11 (44) | 6 (40) | 5 (27) | 7 (32) | 4 (18) |
| Type III | 2 (20) | 0 (0) | 4 (18) | 1 (10) | 3 (12) | 1 (7) | 1 (6) | 4 (18) | 4 (18) |
| Type IV | 0 (0) | 0 (0) | 0 (0) | 0 (0) | 0 (0) | 0 (0) | 1 (6) | 0 (0) | 1 (5) |

**Supplementary Table 4**. Distribution of OFC sulcogyral patterns and associated SOGS scores as a function of left/right hemisphere combination.

|  | Sulcogyral patterns | | Left | | | |
| --- | --- | --- | --- | --- | --- | --- |
|  |  |  | I | II | III | IV |
| Pathological gamblers | Right | I | N=33  (10.00±4.51) | N=32  (10.13±4.01) | N=8  (9.80±4.21) | N=2  (9.50±2.12) |
|  |  | II | N=28  (10.58±3.65) | N=37  (10.15±4.71) | N=5  (9.00±4.85) | N=2  (11.50±4.95) |
|  |  | III | N=3  (12.50±2.12) | N=5  (9.60±2.70) | N=5  (13.75±3.30) | N=0 |
|  |  | IV | N=1  (14.00) | N=2  (10.50±0.71) | N=1  (4.00) | N=1  (17.00) |
|  | Sulcogyral patterns | | Left | | | |
|  |  |  | I | II | III | IV |
| Healthy controls | Right | I | N=38  (0.41±0.82) | N=29  (0.62±1.13) | N=16  (0.27±1.03) | N=1  (0.00) |
|  |  | II | N=26  (0.48±0.95) | N=20  (0.33±0.49) | N=5  (0.50±1.00) | N=2  (0.00±0.00) |
|  |  | III | N=9  (0.44±1.01) | N=6  (0.33±0.82) | N=5  (0.00±0.00) | N=0 |
|  |  | IV | N=1  (0.00) | N=0 | N=1  (0.00) | N=0 |

*The numbers outside the brackets represent the numbers of participants for each left-right combination of the four sulcogyral patterns, while the numbers in brackets represent the mean ± standard deviation of SOGS scores.*
